# The efficacy of a virtual reality exposure therapy treatment for fear of flying: A retrospective study

**DOI:** 10.1101/2020.05.27.118695

**Authors:** Amihai Gottlieb, Glen M. Doniger, Yara Hussein, Shlomo Noy, Meir Plotnik

## Abstract

**Background:** Fear of flying (FoF) is an anxiety disorder classified as a phobia. Its prevalence is estimated at 10–40% in the industrialized world, and it is accompanied by severe economic, social, vocational and emotional consequences. In recent years, virtual reality-based exposure therapy (VRET) for FoF has been introduced. One such FoF-VRET is offered as a paid clinical service at the Center of Advanced Technologies in Rehabilitation (CATR), Sheba Medical Center, Israel. Positive long-term efficacy of FoF-VRET has been found in several studies. However, these studies are limited by relatively small, non-representative samples and a lack of comparative pre/post functional efficacy outcome measures.

**Methods:** To address these limitations, we conducted a retrospective survey of self-referred individuals treated with FoF-VRET at CATR over the previous four years. The aim of the present study was to evaluate the efficacy of our FoF-VRET in this representative real-world sample. Of 274 individuals who received the treatment, 214 met inclusion/criteria, and 103 agreed to participate. The survey focused mainly on collecting information regarding flight activity before and after treatment. The primary outcome measures were: (1) number of flights per month (FpM); (2) number of flight hours per month (FHpM). For each participant, these outcomes were computed for the post-treatment period (≥6 months after FoF-VRET) and the corresponding pre-treatment period.

**Results:** FpM (mean±SD) increased from .05±.07 to .16±.07 flights (*p*<.0001). FHpM rose from.22±.41 to .80±.86 hours per month (*p*<.0001).

**Conclusions:** These results are indicative of FoF-VRET treatment efficacy. Future studies should evaluate long-term maintenance of the treatment effect and thus identify the optimal frequency for delivery of periodic booster treatments.

## I. Introduction

Airplanes are the safest, most common way to travel in the industrialized world (1, 2). Fear of flying (FoF) is a common anxiety disorder in western countries, and its prevalence is estimated at 10–40% (3–5). Among those who suffer from FoF, 14% have never flown on an airplane, 6% have flown and say they will not fly again, and 10% have flown and say they will fly again only if there is no other choice (6). FoF may be secondary to phobias related to environmental conditions (e.g., altitude, severe weather) or situational phobias (e.g., claustrophobia), and may be comorbid to panic attacks and generalized anxiety disorder (7). Physiological and psychological anxiety symptoms of FoF may include panic attacks, fear, muscle tension, sweating, shortness of breath, heart palpitations, nausea and dizziness (8). The costs of FoF for affected individuals, their families and society are substantial. FOF sufferers tend to avoid flying entirely, which may have serious social, vocational and emotional consequences (9). Societally, FoF results in significant cost to the airlines and incalculably reduced productivity and opportunity (9).

Several pharmacological treatments exist for FoF including anti-anxiety medications like benzodiazepines (10). Other treatments are psychological interventions like exposure therapy (also called systematic desensitization) (11). The most common treatment for FoF is cognitive behavioral therapy (CBT) (12), which focuses on creating neutral memories to replace the panic-inducing ones, and may include relaxation techniques, psychoeducation and exposure therapy. In exposure therapy, the FoF sufferer is exposed to the source of anxiety in a controlled manner, and this approach is considered the most efficient treatment for FoF (13, 14). Exposure therapy for FoF might involve simulating a flight or exposure to a stationary plane.

Over the last decade, virtual reality (VR) has become a viable method for administering exposure therapy for anxiety disorders. For example, several VR-based exposure treatments for post-traumatic stress disorder have been proposed [for review see (15)]. As applied to FoF, virtual reality exposure therapy (VRET) involves creating a virtual airplane environment that simulates various aspects of flying using dynamic visual, auditory and motion stimuli (7). Unlike exposure therapy using a real flight, this VR-based method allows the therapist to systematically control the level of the exposure intensity to a variety of elements (16). Notably, VRET for FoF (FoF-VRET) is most often implemented with a VR head mount display (e.g., (17, 18)) and thus lacks the ability to simulate motion. Large-scale VR systems that incorporate motion can be used to address this limitation and better simulate the flight experience.

There are few reports evaluating the clinical efficacy of FoF-VRET [e.g. (13, 17–22)]. In a recent meta-analysis of 11 randomized trials, Cardos and colleagues found FoF-VRET to be superior to control/standard FoF treatments (23). Only a few randomized trials have assessed efficacy in the months following treatment. For example, Rothbaum and colleagues (2000) (21) reported maintenance or enhancement of self-reported post-treatment improvements after six months for both VRET and standard exposure therapy groups. Further, at six months post-treatment, 79% of VRET participants and 69% of standard exposure therapy participants reported that they had flown (voluntarily) since completing the treatment. In another study, Muhlberger and colleagues (2003) (17) found that 62% of VRET participants reported flying during the 6-month follow-up period. However, Maltby and colleagues (2002) (20) reported that differences between VRET and an attention-placebo group observed immediately following treatment had disappeared after six months. In a randomized controlled trial, Rothbaum and colleagues (2002) (24) found that 92% of VRET and 91% of standard exposure participants had flown one-year post-treatment. Tortella-Felui (2011) (18) found that 66% of VRET participants reported flying during the 1-year follow-up period. Finally, in a long-term follow-up study, Wiederhold (2003) (25) found that 85% of their 30 participants reported flying in the three years after completing several different VRET treatments.

Taken together, sample sizes in these studies were relatively small, and it is apparent that there is great variability (62-92%) in the prevalence of (voluntary) flying in the period following the conclusion of VRET treatment (17, 24). Further, participants in such studies are not considered representative of the general population as they have consented to an experimental treatment and are thus particularly motivated and amenable to the treatment. Most importantly, existing studies lack comparative pre/post functional efficacy outcome measures. To address these issues, better controlled studies with larger, more representative clinical samples are needed.

The Center of Advanced Technologies in Rehabilitation (CATR) at Sheba Medical Center (Ramat Gan, Israel) has developed a FoF-VRET using a large-scale VR system. See Czerniak and colleagues (2016) (7) for a full description of the setup (see also *Methods*). To date (January 2019) more than 274 individuals have been treated.

The aim of the present study was to evaluate the efficacy of our FoF-VRET by retrospectively surveying individuals who received the treatment as a paid clinical service. Our primary objective was to evaluate whether flying habits changed after completion of the treatment.

## II. Methods

### Rationale

Between 2014 and 2018, 274 individuals were self-referred to receive FoF-VRET at CATR. We emailed 214 individuals who had completed the treatment and for whom we had an email address on file. In the email, we asked if they would be willing to participate in a phone survey regarding the FoF treatment they received (see *Procedure* for more details). Among the benefits of this methodology are: (1) reduced bias associated with willingness to participate in experimental research; (2) reduced bias associated with an onsite office interview by a clinician; (3) reduced ‘gratitude effect’ consequent to *pro bono* research participation; participants in the present study paid out-of-pocket to obtain a clinical service.

### Participants

Inclusion criteria – (1) completion of the FoF-VRET treatment regimen at CATR; (2) email address on file (to facilitate emailing of consent at initial contact).

Exclusion criteria – (1) non-responsive to email; (2) refusal to participate; (3) <6 months after treatment completion.

Six months was set as the minimum time from treatment completion to allow a reasonable amount of time for participants to fly and for comparability to the literature (see *Main outcome measures*). Of the 214 individuals we contacted, 103 actually participated. Individuals were excluded for the following reasons: 50 were non-responsive, 53 refused to participate and 8 were questioned <6 months from treatment completion.

### Procedure

First, potential participants were emailed for their consent to participate; those who agreed were then contacted by phone to confirm their informed consent. Next, a structured phone interview was conducted. The interview consisted of three parts:

1. Confirmation of FoF-VRET treatment dates and recording the reason or reasons for self-referral.
2. Information on flight activity for the period following treatment completion and a corresponding period of identical length of time prior to treatment initiation. For each flight, participants reported their destination and flight duration. In addition, participants rated their anxiety level during each flight on a scale from 1 (least anxious) to 7 (most anxious). Only outbound flights were recorded (i.e., flights departing Ben Gurion Airport, Israel). For verification purposes, participants were asked to furnish supporting material, including boarding passes and passport stamps.
3. Questions about the FoF-VRET treatment experience, including whether they underwent other FoF treatments ±1 year before/after the FoF-VRET treatments.

### Main outcome measures

The primary measure of FoF-VRET efficacy was number of flights per month (FpM). The secondary outcome measure was number of flight hours per month (FHpM). For example, a participant interviewed 18 months after VRET completion reported the following flight information: to New York (11h) in month +2, to London (5.5h) in month +7, and to Eilat (1h) in month +17. His/her outcome measures were thus FpM=(3/18) and FHpM=(17.5/18). Corresponding pre-treatment measures were calculated from data for the identical period of pre-treatment time. An indirect measure of treatment efficacy was reported anxiety level for pre- and post-treatment flights.

### FoF-VRET treatment

Refer to the Supplementary Material for a brief description of the FoF-VRET treatment (for a full description see *Czerniak, E., Caspi, A., Litvin, M., Amiaz, R., Bahat, Y., Baransi, H., … Plotnik, M. (2016). A novel treatment of fear of flying using a large virtual reality system. Aerospace medicine and human performance, 87(4), 411-416*.).

### Statistical analyses

Non-parametric within-participant analyses (Wilcoxon signed-rank tests) were used to compare pre- and post-treatment FpM, FHpM and anxiety levels. Alpha level was set at *p*<. 05, two-tailed.

## III. Results

### Flight activity before and after FoF-VRET treatment

Participants showed a clear increase in flight activity post-treatment as compared to pre-treatment (Figure 1).

**Figure 1.**
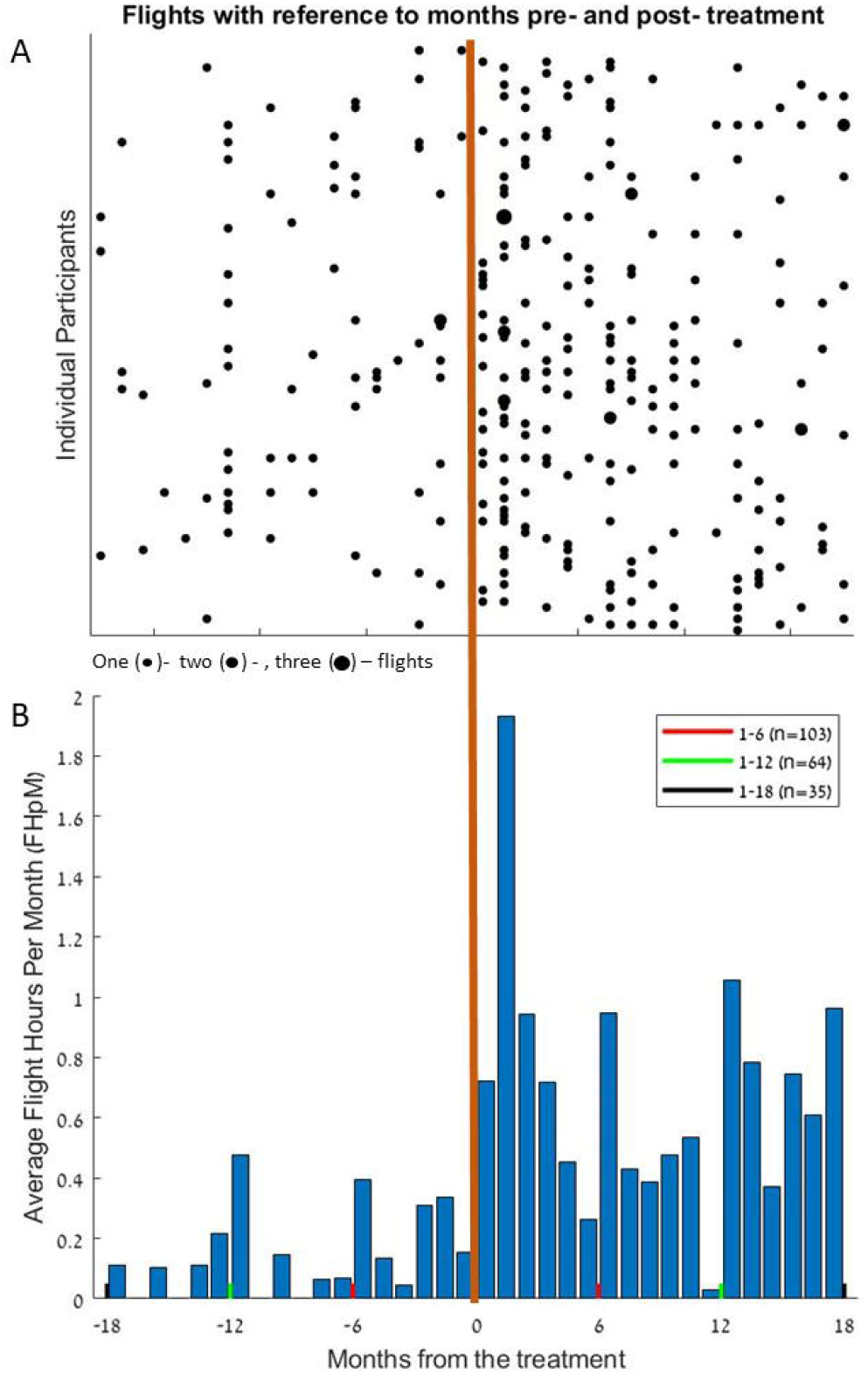
Flight activity 18 months before and after FoF-VRET treatment for individual participants. A. Each point represents at least one flight for the given month (see key below panel). Negative values on the abscissa reflect months pre-treatment, and positive values reflect months post-treatment, vertical orange line represents the month during which the treatment took place. Each horizontal row represent data from one participant. Data from 17 participants who did not fly before or after the treatment (i.e, reciprocal 18 months periods pre- and post-treatment) are not shown, yet these data were included in statistical analyses. Following treatment, mean ± SD FpM increased from .05±.07 to .16±.07 flights (n=103; see also text for non-parametric comparisons). B. Mean flights hours per month (FHpM) across participants. Following treatment, mean FHpM rose from .22±.41 to .80±.86 hours per month. Note that for each participant, pre-treatment data were analyzed for the identical length of time as the post-treatment period at the time of data collection (see text). Thus, for all 103 participants, data was analyzed for 6 months pre/post treatment (red lines), for 64 participants data was analyzed for 12 months pre/post treatment (green lines), and for 35 participants data was analyzed for 18 months pre/post treatment (black lines). Pre-hoc analyses confirmed uniformity of distributions during overlapping periods for all three groups.

Regarding flight activity outcomes before and after treatment, within-participant analyses revealed a significant difference for FpM and FHpM before (FpM: median=0, Interquartile Range (IQR)=.07, FHpM: median=0, IQR=.28) and after (FpM: median=.13, IQR=.18, FHpM: median=.57, IQR=1.04) treatment (Wilcoxon signed-rank tests, *Z*=-6.89, *p*<.0001, *Z*=-6.73, *p*<.0001, respectively).

Figure 2 shows FpM and FHpM across participants in the months before and after treatment.

**Figure 2.**
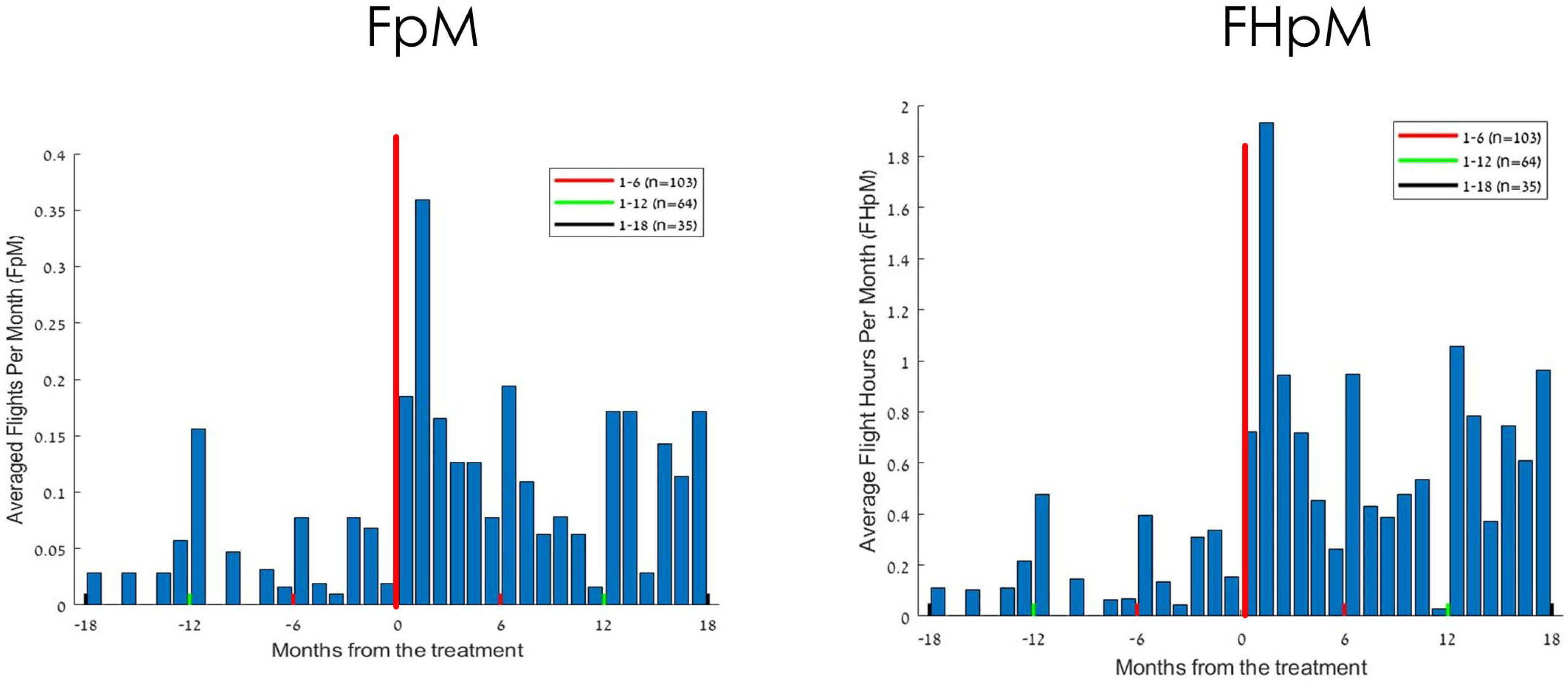
Mean flights per month (FpM; left) and flight hours per month (FHpM; right) across participants. Negative values on the *x*-axis reflect months pre-treatment, and positive values reflect months post-treatment. Following treatment, mean±SD FpM increased from .05±.07 to .16±.07 flights; mean FHpM rose from .22±.41 to .80±.86 hours per month. Note that for each participant, pre-treatment data was analyzed for the identical length of time as the post-treatment period at the time of data collection (see text). Thus for all 103 participants, data was analyzed for 6 months pre/post treatment (red lines), for 64 participants data was analyzed for 12 months pre/post treatment (green lines), and for 35 participants data was analyzed for 18 months pre/post treatment (black lines).

### Anxiety before and after FoF-VRET

Within-participant analyses (Wilcoxon test) revealed a significant difference in self-reported anxiety level for flights before (median=6.0, IQR=2.0) and after (median=4.0, IQR=2.2) the treatment (Wilcoxon signed-rank test, *Z*=-5.16, *p*<.0001). Figure 3 shows anxiety level for flights before and after FoF-VRET.

**Figure 3.**
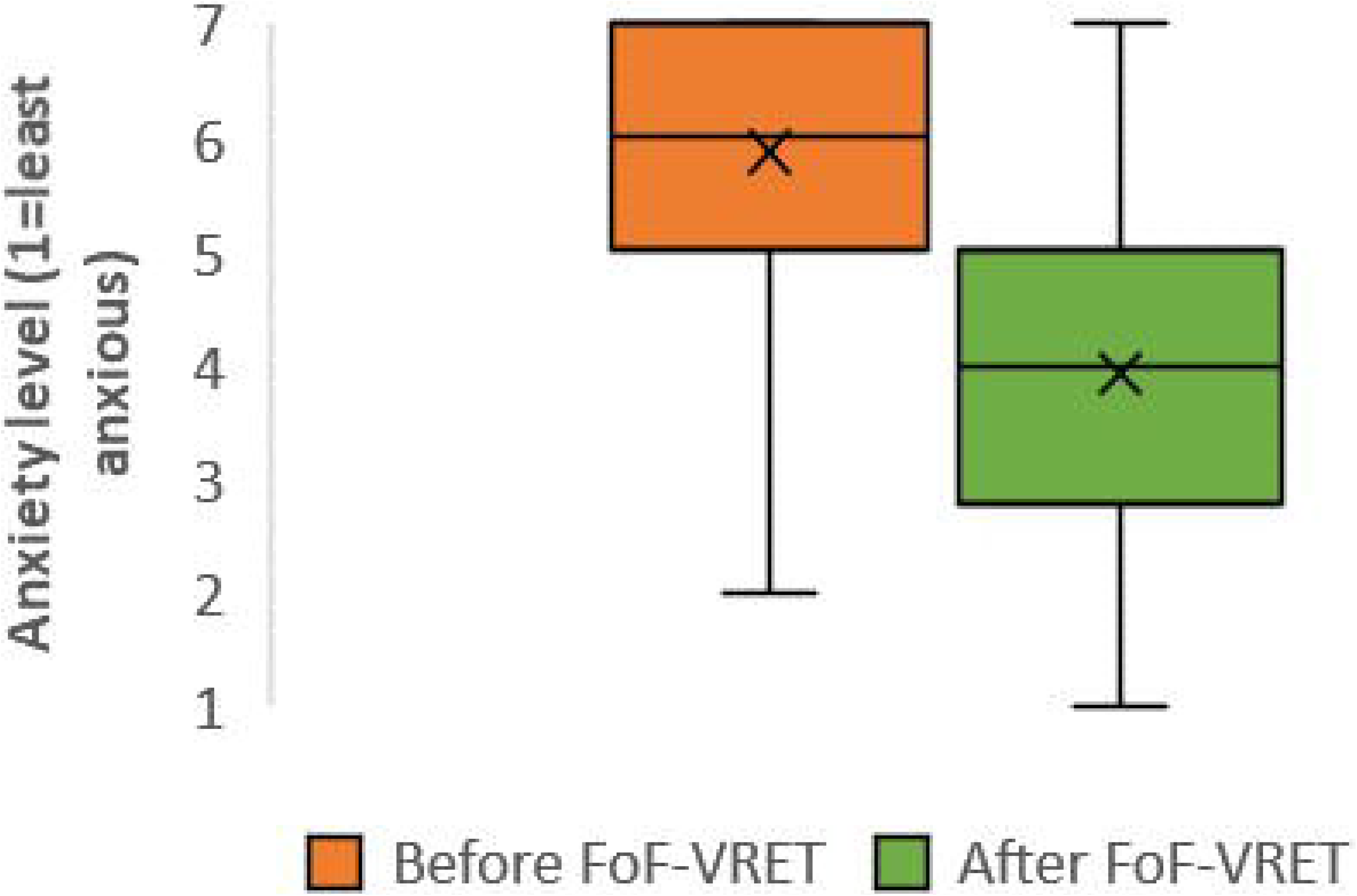
Level of self-reported anxiety across participants for flights before and after FoF-VRET (box plots). The scale ranged from 1 (least anxious) to 7 (most anxious).

### Reasons for seeking FoF-VRET and other FoF treatments

To provide additional clinical background, Figure 4 shows the distribution of reasons for seeking FoF-VRET across participants, as reported during the phone interview.

**Figure 4.**
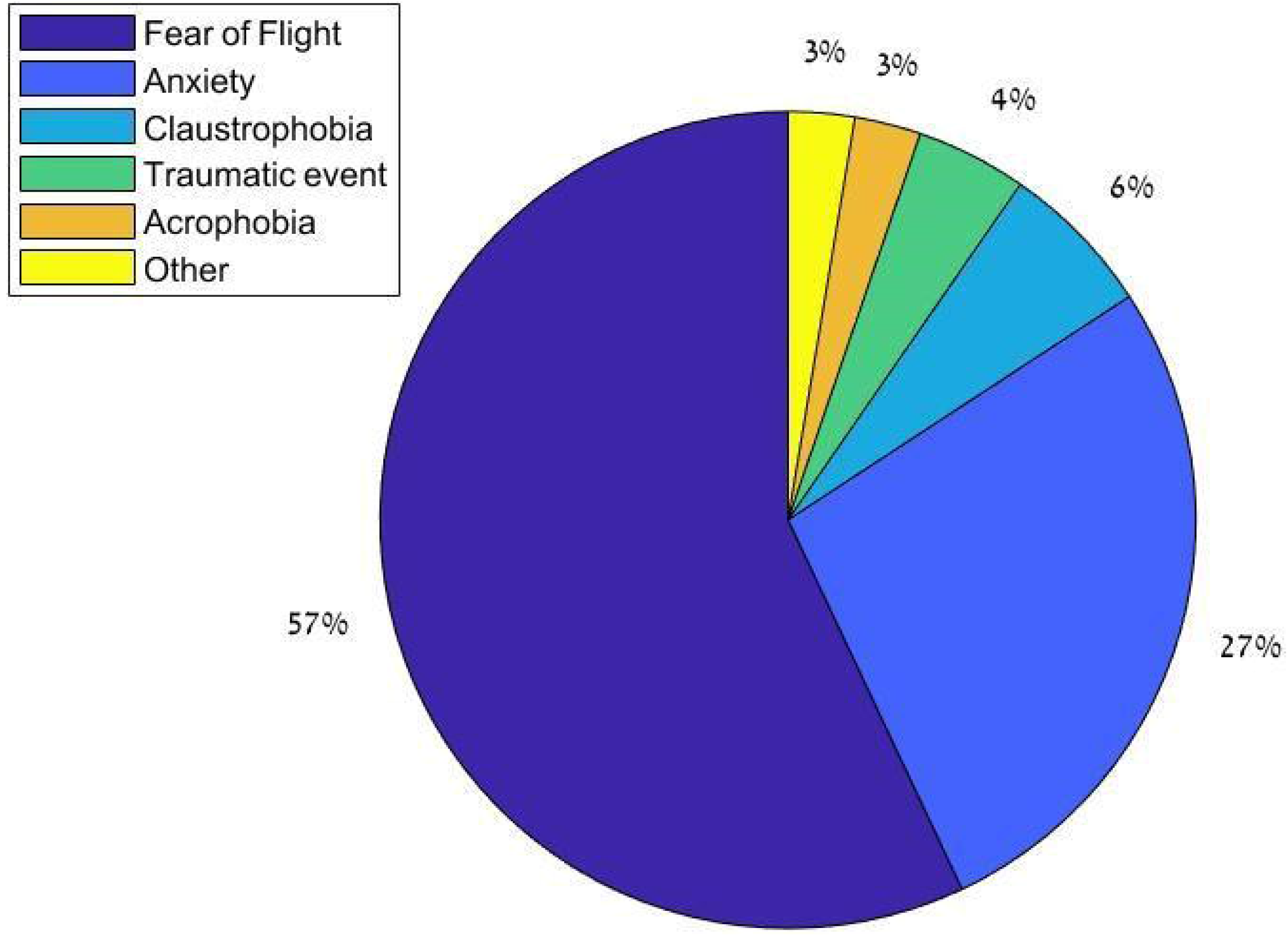
Distribution of reasons for seeking FoF-VRET treatment across participants. Note: For participants who reported more than one reason, all reported reasons are included.

Figure 5 shows the distribution of other treatments for FoF within approx. ±1 year of FoF-VRET reported by participants. The distribution indicates that nearly half (48%) of participants did not engage in any other treatment. Among the other participants, psychological treatment (18%) and FoF workshops (12%) were most common; hypnosis (4%) and CBT (3%) were least common.

**Figure 5.**
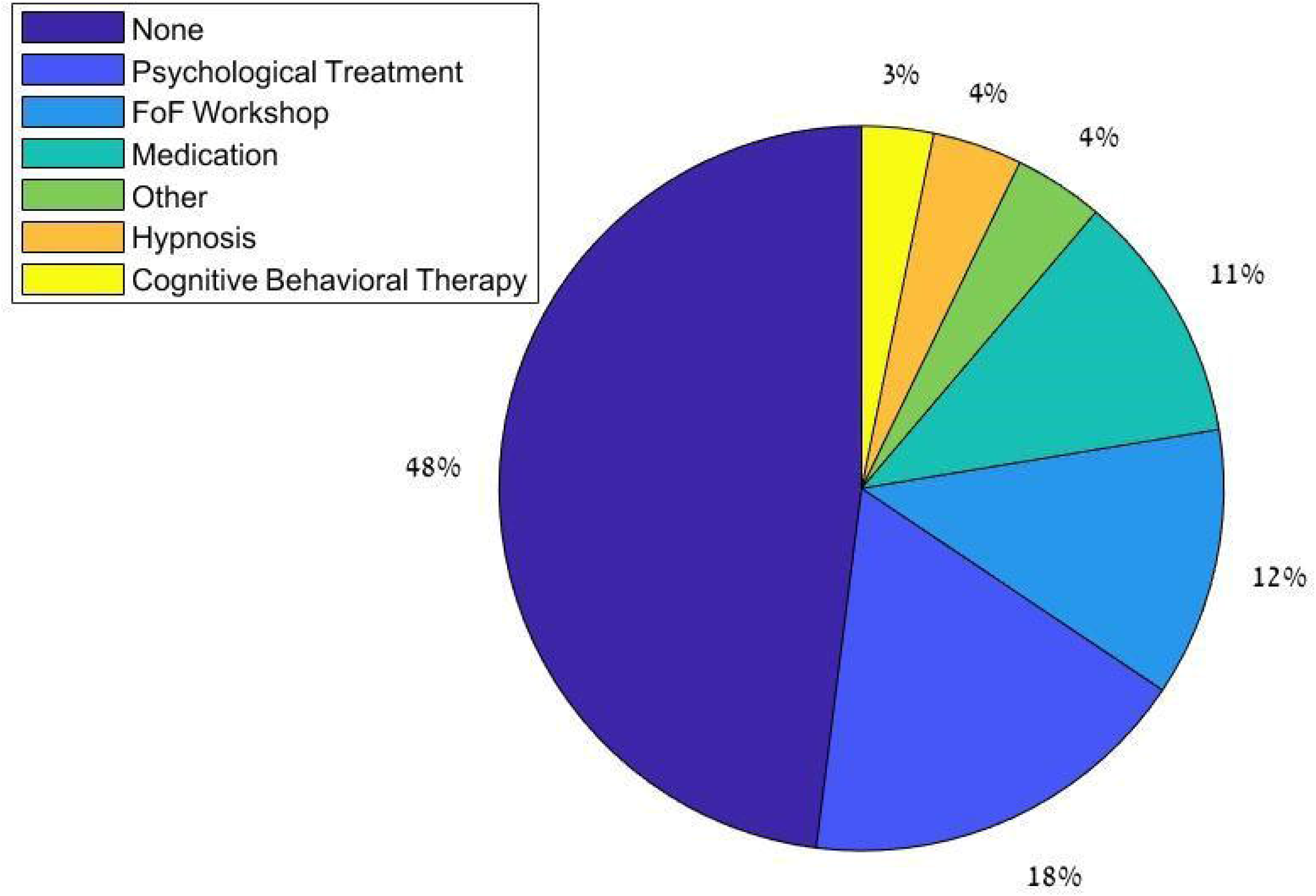
Distribution of other FoF treatments within one year of FoF-VRET. Note: For participants who reported more than one treatment, all reported treatments are included.

To confirm that our findings regarding FoF-VRET efficacy were not unduly affected by the additional treatments, we conducted a post hoc analysis. Briefly, we split participants into ‘no other treatment’ (48%), and ‘other treatment’ groups. For each participant, we computed pre/post change (post-treatment minus pre-treatment; Δ) in FpM and FHpM, respectively. Man-Whitney tests revealed no significant differences between the groups (ΔFpM: *U*=1397.5, *p*=.56, ΔFHpM: *U*=1379.5, *p*=.64), suggesting that other treatments did not appreciably affect the increased flight activity we attribute to FoF-VRET.

## IV. Discussion

This study aimed to determine the efficacy of FoF-VRET treatment using a retrospective follow-up questionnaire conducted over the phone. Our study is novel in that we evaluated individuals who voluntarily paid for and received treatment in our virtual reality center.

This FoF-VRET has several advantages over standard FoF exposure treatments: Firstly, it provides a safe, controlled environment that can be continuously monitored and manipulated by professional therapists and technicians. Secondly, it provides a highly detailed visual, auditory, and motion simulation of an actual flight experience rather than a static airplane. This provides better exposure to the fear-triggering factors, potentially inducing participant responses more similar to those elicited by real air travel. Finally, other measures like heart rate and blood pressure can be recorded during VR exposure therapy to provide therapists with more comprehensive clinical information.

### Efficacy

The current results show that number of flights and flight hours post-treatment significantly increased, reflecting treatment efficacy. Additionally, in-flight anxiety level significantly declined following treatment. These results are in line with other retrospective follow-up studies assessing the efficacy of other virtual reality-based FoF treatments (17, 18, 20, 21, 24, 25). The results of this study corroborate these prior studies and provide new evidence that those who benefited from the treatment continue to fly as long as eighteen months after FoF-VRET treatment initiation.

While previous studies evaluating the efficacy of FoF-VERT used air travel in the post-treatment period (i.e., yes/no) as the sole (binary) outcome (e.g. (24, 25)), the current study introduces additional measures: flight frequency (i.e., number of flights per month) and flight hours per month in the post-treatment period. We believe that with the addition of these measures, we are able to provide better evidence of treatment efficacy, as we show that treated participants no only fly more often, but also that they fly for longer durations. These results suggest that engaging in FoF-VRET leads participants to take flights they would not have been prepared to take prior to treatment.

In a recent meta-analysis of 11 randomized controlled trials, Cardos et al. (23) reported significant overall efficacy of a FoF-VRET intervention (*G*=0.592) and a significant increase in flight activity at follow-up (*G*=0.588), demonstrating the advantage of FoF-VRET treatment over control/traditional FoF treatments. However, their results also reveal the limitations of these trials due to poor study quality and small sample size. The authors suggest that reported effects may have been overestimated as a result of these issues. In contrast, our findings are based on a larger sample size and a more true-to-life (ecological) environment than those of the aforementioned studies.

### Other results

Some of our results elucidate clinical aspects of FoF and its treatment. While most participants reported suffering specifically from FoF (acrophobia), a significant number of participants reported suffering from general anxiety. Furthermore, almost half (48%) of the individuals receiving FoF-VRET treatment reported that they did not engage in any other treatments at least one year prior to treatment, suggesting that half of those suffering from FoF are untreated and may avoid air travel.

### Limitations and future work

The current study is limited in several important ways. Firstly, to maximize sample size, we did not collect data at a fixed length of time from treatment (e.g., one year). Consequently, some adjustments to the data were required (e.g., standardizing the primary outcome measures to permit within-subject statistical comparisons). Secondly, the attrition rate was relatively high (51.8%), which may have affected the results. Although this level of attrition was higher than in other retrospective follow-up studies (13% in Rothbaum et al. (24), 29.3% in Tortilla-Feliu et al. (18), 10% in Wiederhold et al. (25) and 10% in Muhlberger et al. (17)), a higher attrition rate may be expected for participants solicited to participate in a phone survey following receipt of a clinical treatment they paid for as compared to participants volunteering in research studies.

### Conclusions

Current results are indicative of FoF-VRET treatment efficacy. Air travel is an integral part of modern life in the industrialized world, and its prevalence is expected to grow as airfares continue to decrease and global economics entails more business travel (1). We can therefore expect a heightened awareness of FoF and an increase in referrals for suitable treatments including VRETs. Future studies should evaluate long-term maintenance of the treatment effect and consequently identify the ideal frequency for delivery of subsequent booster treatments.

## List of abbreviations

CATR: Center of Advanced Technologies in Rehabilitation
CBT: Cognitive Behavioral Therapy
FHpM: Flight Hours per Month
FoF: Fear of Flying
Fpm: Flight per Month
IQR: Interquartile Range
VR: Virtual Reality
VRET: Virtual Reality Exposure Treatment

## Acknowledgments

We would like to thank Ms. Yarden David and Ms. Yael Menkes, for performing some of the phone surveys.

