## Supplementary Material for "The efficacy of a virtual reality exposure therapy treatment for fear of flying: A retrospective study"

##### Description of FoF-VRET Treatment

The following brief description of the treatment is taken with permission from Cherniak et al. (2016).

**“**We used the Computer Assisted Rehabilitation Environment (CAREN; Motek Medical ©, Amsterdam, the Netherlands) high-end system (Fig. 1A). The virtual visual scenery depicts the interior of an aircraft, where several passengers are seated (Fig. 1B). The virtual outside world can be viewed constantly through the window, e.g., on-the-ground airfield sceneries (e.g., terminal, runways). The complete ground scenery is seen in Fig. 1C. The virtual areal path was designed as a circular route, starting at the airport, over fields and countryside, above the clouds, into an urban area, and back to the airport (Fig. 2). The treated patient was seated in the middle of a motion platform (Fig. 3A). The flight simulation lasted 35 min (6 min of ground pre-takeoff, 24 min for takeoff and airborne, and 5 min of ground post-landing) and included the following stages: taxiing, takeoff (Fig. 3B), cruise (including turns), and landing. The inclinations of the platform (2D, pitch, roll) were congruent with the rotations of the visual scenery. The platforms were also linearly accelerated (3D), e.g., for 'vertical drops' (simulating air turbulences). Corresponding views can be seen through windows either on the same side or on the opposite side of the aircraft based on its virtual location and orientation. A portion of the aircraft 's wing is constantly displayed. The auditory exposure includes jet engine sound, i.e., the engines starting and increasing power during takeoff, as well as vocal announcements (i.e., made by the flight crew). Throughout the flight simulation, the operator can introduce several exposure levels of simulated unexpected events including: 1) turbulences, 2) fire and smoke (Fig. 3C), 3) night flight (not applied in the present cases), and 4) smooth/ rough landing. In the first session the therapist (a clinical psychologist) interviewed the patient regarding his FoF and other specific phobias. The treatment plan was then scheduled accordingly. During the interview the therapist provided the patients with the safety aspects of flights, including emergency procedures and the actions taken by the pilots during technical malfunctions. Next came three consecutive (once a week) VRET sessions in the presence and with the guidance of the therapist. Prior to each VRET, there was a short CBT session confronting the emotions-thoughts-behavior aspects of the patient’s core fear, cognitive restructuring, and psychoeducation for providing anxiety-coping skills, as well as breathing and relaxation techniques**”**.

Fig. 3. A) The passenger aircraft seat was located in the middle of the platform (a standard chair was used). B) The patient is seated for takeoff. The platform is inclined backward, and the aircraft’s wing can be seen through the windows. C) A simulated incidence of one engine catching fire and the smoke following it. *(from Cherniak et al., 2016, with permission).*

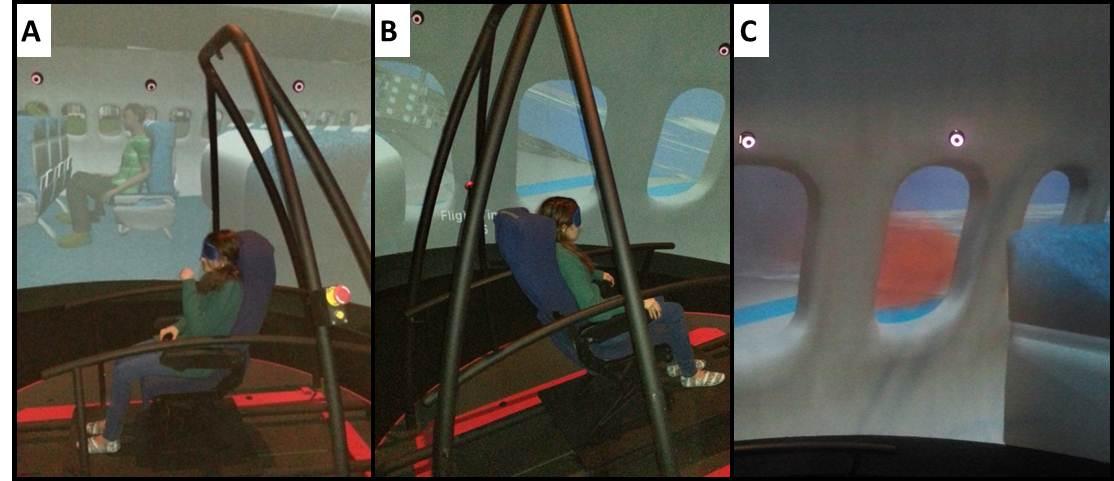

Fig. 1. Virtual reality treatment apparatus, scenery, and flight simulation. A) A schematic drawing of the CAREN high-end system. The system consists of a motion platform (3 m in diameter), which is placed within a dome shape construction. On this interior surface visual scenery is presented using eight projectors which provide a 360° display. Although the projection is not stereoscopic (unlike the case of polarized goggles or head-mounted display), its projection on a spherical screen fully immerses the patient and allows the perception of depth. A surround sound system provides auditory stimuli congruent with the scenery. The platform can be moved (rotations and translations — 6 degrees of freedom) in various velocities and inclinations (image courtesy of Motekforcelink©). B) General impression of the visual scenery. The viewpoint is from a passenger sitting toward the rear end of the aircraft. C) A depiction of the complete sequence of environmental scenes viewed through the window by the patient when the aircraft is on the ground (pulling out from the gate and taxiing). *(from Cherniak et al., 2016, with permission).*

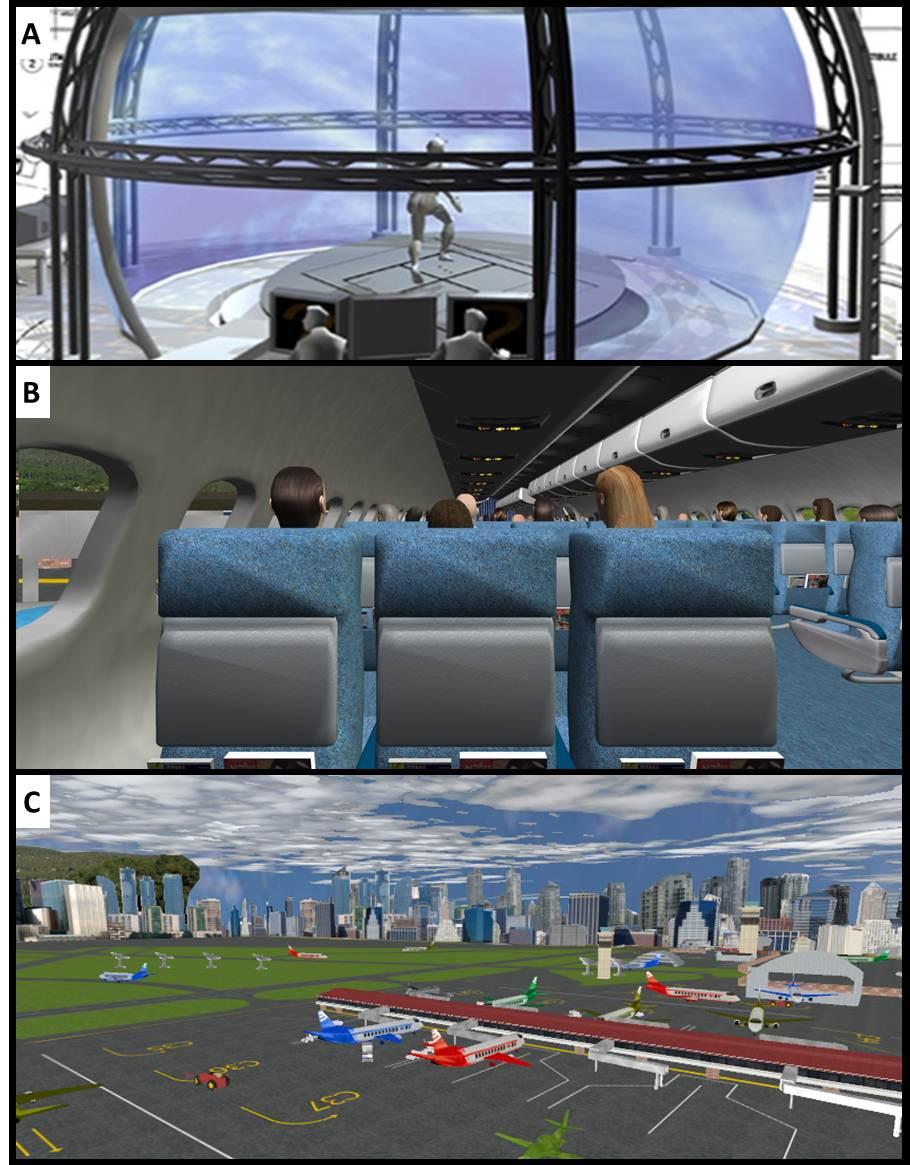

Fig. 2. Simulated flight route and takeoff. The visual illusion of flying was obtained by the creation of a 3D environment (using SoftImage XSI) and then its integration with corresponding sound, platform movements, virtual camera maneuvers, flight conditions, and malfunctions using D-Flow Software (Motekforcelink©). In brief, the virtual plane takes off, flies at 720 km/h, and approaches landing at the speed of 400 km/h — a circular route of 229 km *(from Cherniak et al., 2016, with permission).*

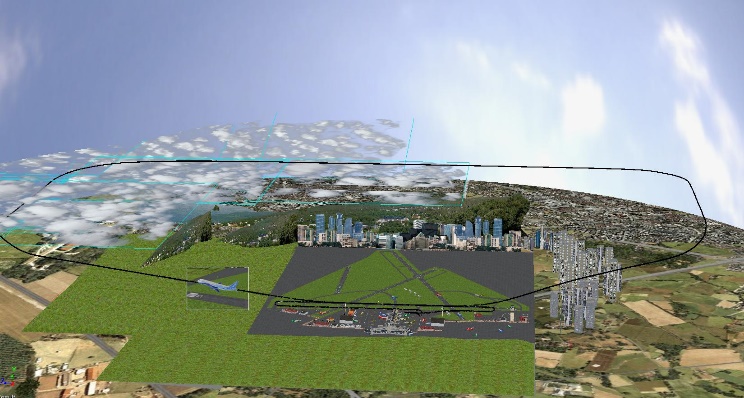
